# IQGAP3 Overexpression Correlates with Poor Prognosis and Radiation Therapy Resistance in Breast Cancer

**DOI:** 10.1101/346163

**Authors:** Xin Hua, Zhi-Qing Long, Wen-Wen Zhang, Chao Lin, Xiao-Qing Sun, Wen Wen, Zi-Jian Lu, Na Guo, Zhen-Yu He, Li Bing-Song, Ling Guo, Huan-Xin Lin

**Affiliations:** State Key Laboratory of Oncology in South China, Collaborative Innovation Center for Cancer Medicine, Department of Radiotherapy, Cancer Center, Sun Yat-sen University, No. 651, Dongfeng Road East, Guangzhou 510060, China; State Key Laboratory of Oncology in South China, Collaborative Innovation Center for Cancer Medicine, Department of Nasopharyngeal Carcinoma, Cancer Center, Sun Yat-sen University, No. 651, Dongfeng Road East, Guangzhou 510060, China

**Keywords:** breast cancer, IQGAP3, prognosis, radiation therapy, resistance

## Abstract

Background: IQ motif-containing GTPase activating protein 3 (IQGAP3), the latest found protein of IQGAP family, may act as a crucial factor in the process of cancer development and progression; however, its clinical value in breast cancer remains unestablished so far. Our team explored the correlation between IQGAP3 expression profile and the clinicopathological features in breast cancer. Methods: IQGAP3 levels in breast cancer cell lines and tumor tissues were detected by real-time PCR and western blotting and compared to the normal control groups. Protein expression of IQGAP3 was evaluated immunohistochemically in specimens (archived paraffin embedded) of 257 breast cancer patients. We also analyze the association between IQGAP3 expression and the clinical characters and prognosis. The relationship between IQGAP3 expression and sensitivity to radiation therapy was determined by subgroup analysis. Results: There was significant upregulation of IQGAP3 in breast cancer cell lines and human tumor tissues at both the mRNA and protein level compared to the normal ones. In addition, 110/257 (42.8%) of archived paraffin embedded breast cancer specimens had high protein expression of IQGAP3. High expression of IQGAP3 was significantly related to clinical stage (P=0.001), T category (P=0.002), N category (P=0.001), locoregional recurrence(P=0.002), distant metastasis (P=0.001), and vital status (P=0.001). Univariate and multivariate statistical analysis showed that IQGAP3 was an independent prognostic factor of the whole cohort breast cancer patients (P=0.003, P=0.001). Subgroup analysis revealed IQGAP3 expression correlates with radiation therapy resistance and was also an independent predictor for radiation therapy outcome. Conclusions: Our findings suggest that high IQGAP3 expression predicts poor prognosis and radiation therapy resistance in breast cancer. In addition, IQGAP3 may be a reliable novel biomarker to provide personalized prognostication and identify patients who can profit from more aggressive RT regimen for improving the survival of breast cancer patients.

## 1. Introduction

Breast cancer is the most frequent malignancy in women and the second leading cause of cancer-related deaths worldwide[1]. Radiation therapy (RT) is an indispensable part of the systemic therapeutic regimen of breast cancer. Despite great progress in RT over recent years, locoregional recurrence and distant metastasis after RT remain key problems decreasing the survival of breast cancer patients [2]. Locoregional recurrence results from the presence or evolution of radioresistant tumor cells for which standard fractionated RT doses are sublethal[3]. Currently, there is a dearth of clinically available predictive biomarkers to indicate the optimal radiation dosing in breast cancer, which leads to suboptimal treatment of these patients[4]. Therefore, further researches to identify novel biomarkers associated with disease prognosis and RT sensitivity to provide personal accurate prediction, as well as to optimize RT treatment plans, and potential therapeutic targets in breast cancer are urgently required.

IQ motif-containing GTPase activating protein 3(IQGAP3), the latest found protein of the IQGAP family, is an evolutionarily conserved GTPase-activating protein[5] and a hotspot for gene amplification in tumors. IQGAPs comprise five conserved domains: an IQ domain with four IQ motifs (IQ), a poly-proline protein-protein domain (WW), a calponin homology domain (CHD), a RasGAP-related domain (GRD), and carboxy-terminal domain (RasGAPC)[6]. Multiple proteins interact with these domains to regulate diverse cellular processes, including cell migration, cytokinesis, vesicle trafficking, cell proliferation, intracellular signalling, and cytoskeletal dynamics[7].

Overexpression of IQGAP3 has been observed in lung, liver, pancreatic and gastric cancer[8-11]. Recently, IQGAP3 expression was found to be related to clinical stage and was an independent prognostic classifier of gastric cancer patients: IQGAP3 knockdown reduced both the number and size of the spheres formed by the gastric cancer cell line MKN-74 and inhibited the phosphorylation of Akt and Erk1/2[11]. In hepatocellular carcinoma, IQGAP3 was reported to function as an important regulator of epithelial-mesenchymal transition (EMT) and metastasis by activating the transforming growth factor (TGF)-β signaling pathway[9]. Regarding breast cancer, IQGAP3 knockdown by RNA interference inhibited cell proliferation and invasion in two breast carcinoma cell-lines[12]. Although these few reports given clues on the expression of IQGAP3 in several kinds of cancer and its role in tumor development, the correlation between IQGAP3 expression and prognosis or RT sensitivity in breast cancer was still unclear.

Therefore, we inspected the IQGAP3 expression pattern in breast cancer cell lines and patient tissues and in normal control cell lines and tissues. We also analyzed the association of IQGAP3 protein expression with the survival outcomes and RT sensitivity in breast cancer patient cases.

## 2. Results

### 2.1 IQGAP3 expression increased in breast cancer cell lines and breast cancer tissues compared with normal controls

We found IQGAP3 mRNA level was overexpressed in tumor samples compared with normal tissues by analyzing the publically-available microarray TCGA data for breast cancer (Fig.1A). A significant higher expression of IQGAP3 of both mRNA and protein level were observed in the all 12 breast cancer cell lines compared with the control cell line of MCF-10A (Fig.2A).

We used quantitative real-time PCR and Western blotting to evaluate IQGAP3 expression at mRNA and protein levels in tumor samples and the normal control tissues from six different patients. Breast cancer tissues had obviously higher IQGAP3 mRNA and protein levels expression compared with normal breast tissues (Fig.2B). Therefore, the above results prove IQGAP3 is overexpressed in breast cancer.

**Figure1.**
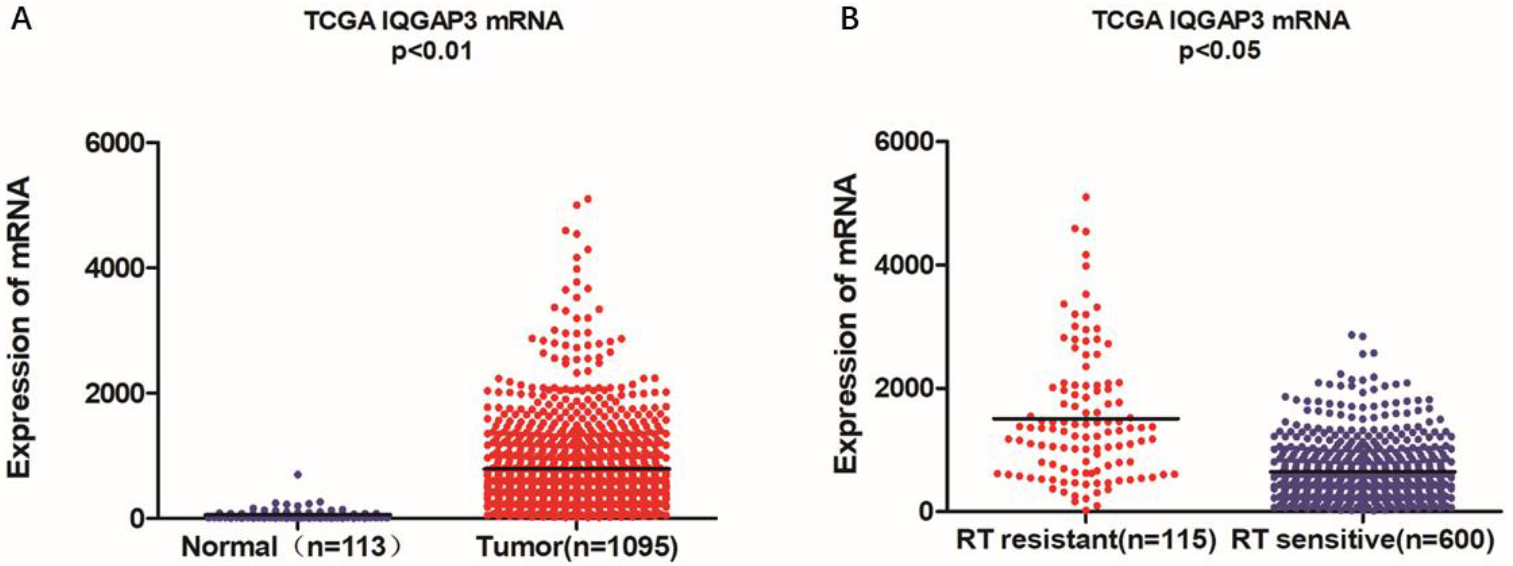
Microarray data reveals IQGAP3 is upregulated in breat cancer patients and in cases that are resistant to radiotherapy (RT). (A) Expression of IQGAP3 in TCGA (breast invasive carcinoma) tumor and normal tissue data (Mann-Whitney test; P < 0.01). (B) Expression of IQGAP3 in TCGA (breast invasive carcinoma) including RT resistant and RT sensitive cases (Mann-Whitney test; P < 0.05).

**Figure 2.**
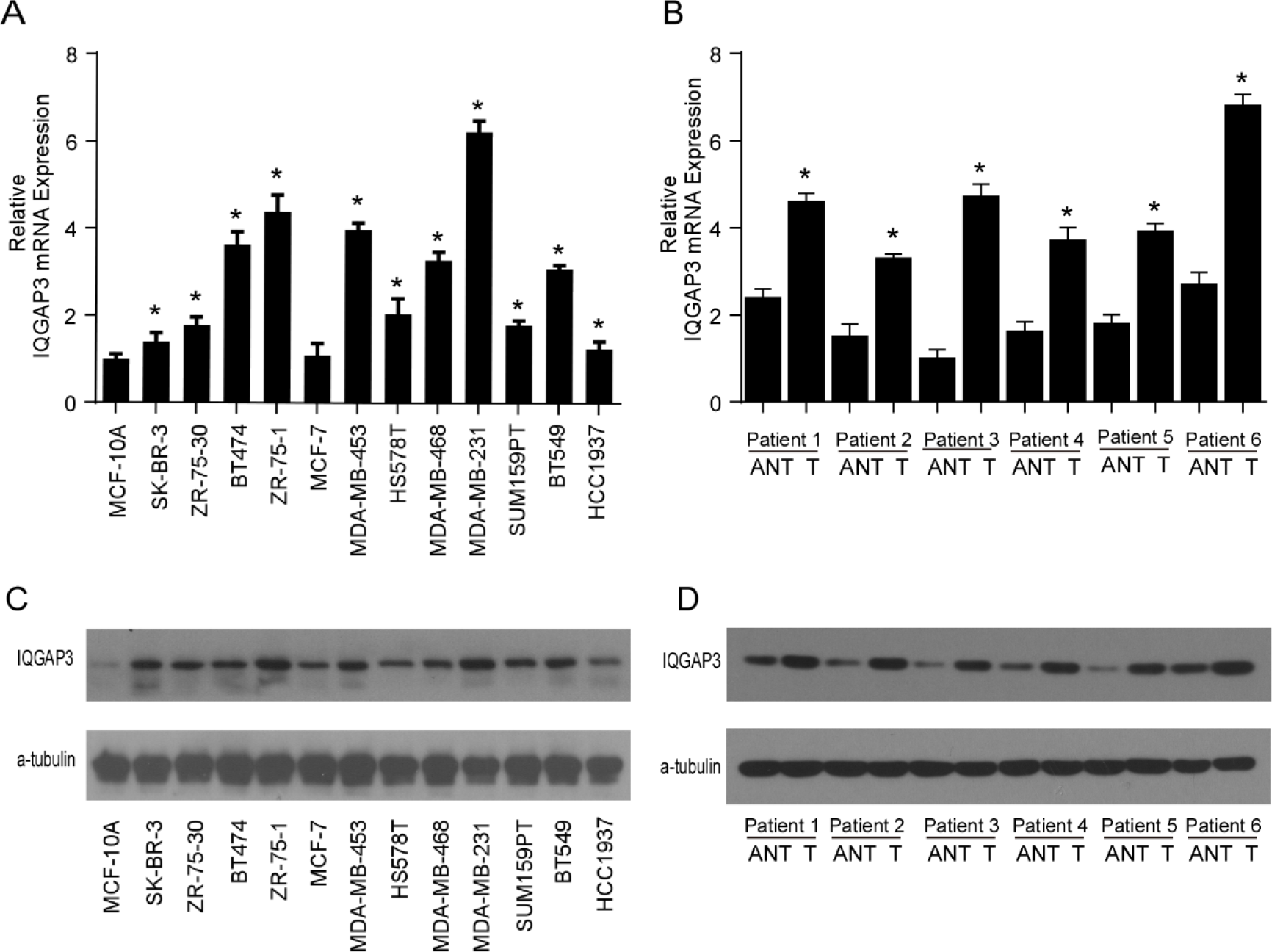
IQGAP3 is upregulated in breast cancer cell lines and tissues. (A) Reverse transcription (RT)-PCR and real-time PCR analysis of IQGAP3 mRNA expression in MCF-10A immortalized breast epithelial cells and twelve cultured breast cancer cell lines. GAPDH was used as a loading control. * P<0.05. (B) real-time PCR analysis of IQGAP3 mRNA expression in six paired breast cancer tumor tissues and normal adjacent tissues(ANT); GAPDH was used as a loading control. * P <0.05. (C) Western blotting analysis of IQGAP3 protein expression in MCF-10A immortalized breast epithelial cells and twelve cultured breast cancer cell lines. α-tubulin was used as a loading control. (D) Western blotting analysis of IQGAP3 protein expression in six paired breast cancer tumor tissues and normal adjacent tissues(ANT); α-tubulin was used as a loading control.

### 2.2. IQGAP3 overexpression was correlated with the clinicopathological features of breast cancer

We investigated the association between IQGAP3 overexpression and the clinicopathological features in the 257 cases of breast cancer specimens by immunohistochemistry. Patients distribution of stage was as follows: stage I (n=21, 8.2%), stage II (n=32, 12.5%), and stage III (n=204, 79.4%). A total of 110 samples (42.8%) demonstrated a higher IQGAP3 protein level expression (staining was strongly positive), and a lower expression (staining was weakly positive or negative) in 147 samples (57.2%, Table 1). Positive IQGAP3 staining was observed mainly in the cancer cell nuclei (Fig.3). IQGAP3 overexpression had a significant correlation with the following characteristics: clinical stage (P=0.001), T category (P=0.010), N category (P=0.001), distant metastasis (P=0.001), locoregional recurrence (P=0.002), and vital status (P=0.001; Table 1).

**Figure 3.**
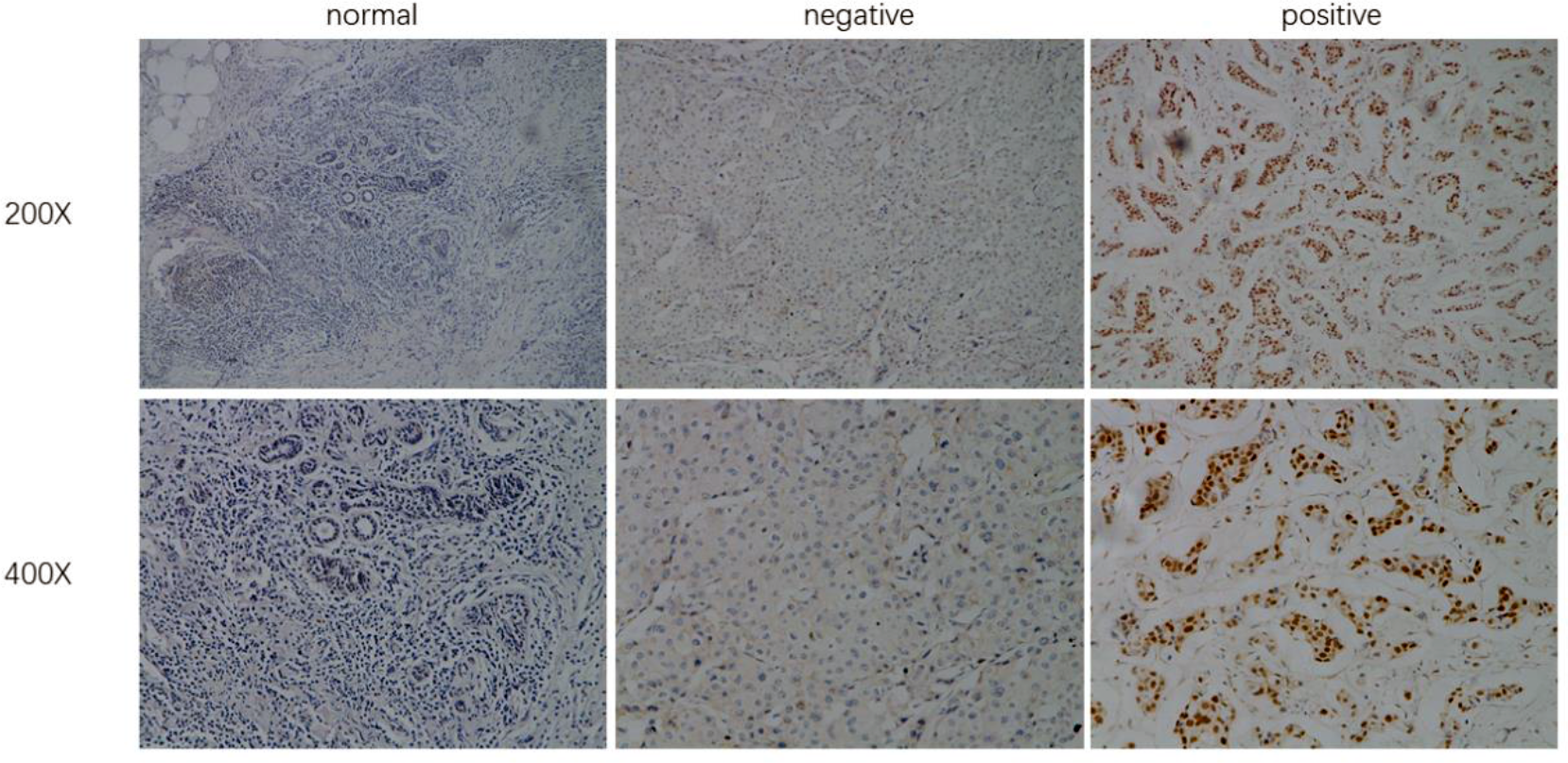
Immunohistochemical detection of IQGAP3 expression in paraffin-embedded breast cancer tissues. Representative images of immunohistochemical staining for IQGAP3 in normal breast tissues(controls) and breast tumor tissues.

**Table 1.**
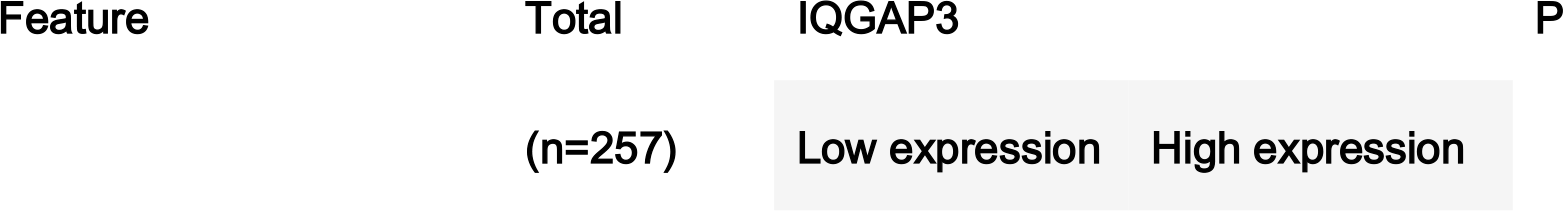
Association between IQGAP3 expression and clinicopathological features of breast cancer (n=257).

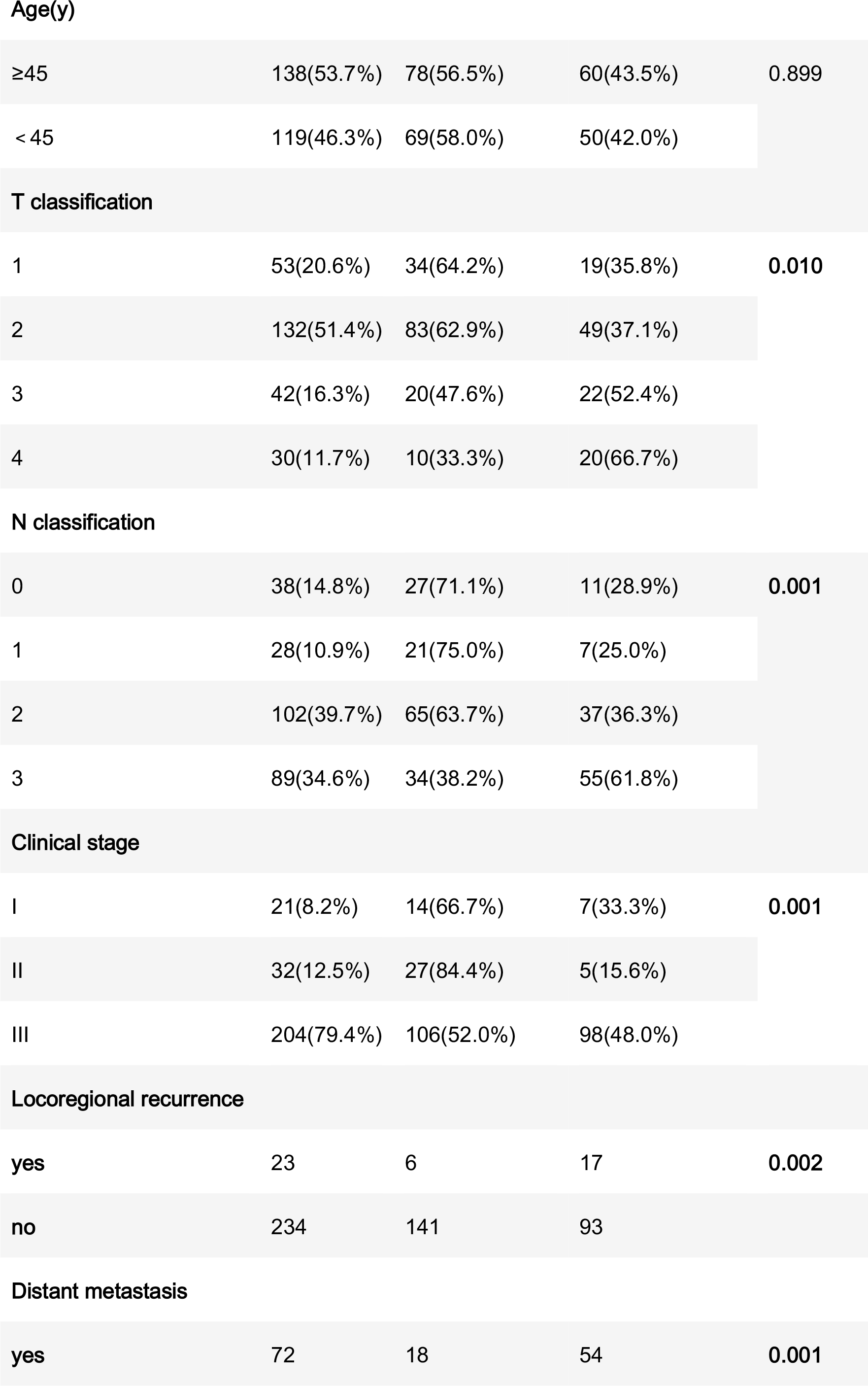

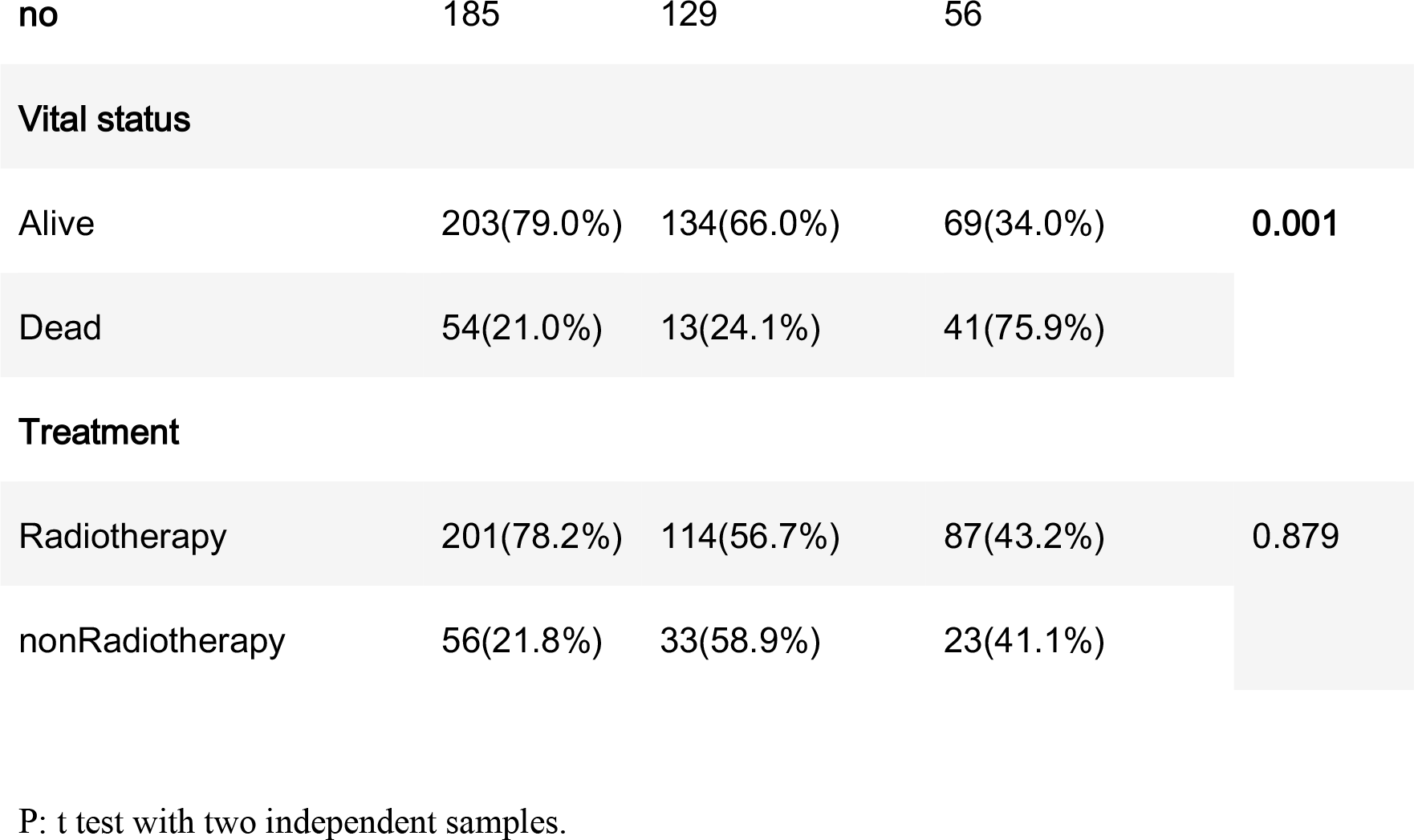

**Age(y)**
|  |  |  |  |  |
| --- | --- | --- | --- | --- |
| ≥45 | 138(53.7%) | 78(56.5%) | 60(43.5%) | 0.899 |
| < 45 | 119(46.3%) | 69(58.0%) | 50(42.0%) |  |
| **T classification** |  |  |  |  |
| 1 | 53(20.6%) | 34(64.2%) | 19(35.8%) | 0.010 |
| 2 | 132(51.4%) | 83(62.9%) | 49(37.1%) |  |
| 3 | 42(16.3%) | 20(47.6%) | 22(52.4%) |  |
| 4 | 30(11.7%) | 10(33.3%) | 20(66.7%) |  |
| **N classification** |  |  |  |  |
| 0 | 38(14.8%) | 27(71.1%) | 11(28.9%) | 0.001 |
| 1 | 28(10.9%) | 21(75.0%) | 7(25.0%) |  |
| 2 | 102(39.7%) | 65(63.7%) | 37(36.3%) |  |
| 3 | 89(34.6%) | 34(38.2%) | 55(61.8%) |  |
| **Clinical stage** |  |  |  |  |
| I | 21(8.2%) | 14(66.7%) | 7(33.3%) | 0.001 |
| II | 32(12.5%) | 27(84.4%) | 5(15.6%) |  |
| III | 204(79.4%) | 106(52.0%) | 98(48.0%) |  |
| **Locoregional recurrence** |  |  |  |  |
| yes | 23 | 6 | 17 | 0.002 |
| no | 234 | 141 | 93 |  |
| **Distant metastasis** |  |  |  |  |
| yes | 72 | 18 | 54 | 0.001 |

| no | 185 | 129 | 56 |  |
| --- | --- | --- | --- | --- |
| Vital status |  |  |  |  |
| Alive | 203(79.0%) | 134(66.0%) | 69(34.0%) | 0.001 |
| Dead | 54(21.0%) | 13(24.1%) | 41(75.9%) |  |
| Treatment |  |  |  |  |
| Radiotherapy | 201(78.2%) | 114(56.7%) | 87(43.2%) | 0.879 |
| nonRadiotherapy | 56(21.8%) | 33(58.9%) | 23(41.1%) |  |

P: t test with two independent samples.

### 2.3. High IQGAP3 expression had significant association with poor prognosis in breast cancer

Overall, of the entire cohort, the 5-year overall survival (OS), locoregional recurrence-free survival (LRFS) and distant metastasis-free survival (DMFS) rates were as follows: 76.9%, 90.7% and 71.9%, respectively. The cumulative 5-year OS, LRFS, and DMFS rates for patients with high IQGAP3 expression were 58.3%, 83.2%, and 50.8%, respectively, compared with 89.9%, 96.1%, and 88.2%, respectively, for patients with low IQGAP3 expression (P=0.001; Fig.4A, P=0.001; Fig.4B and P=0.001; Fig.4C).

Furthermore, the prognostic predictive value of IQGAP3 overexpression in the RT subgroup (n=201) was evaluated. Significant association was found between IQGAP3 high expression and poor OS, LRFS and DMFS in patients who had undergone RT (P=0.001; Fig.4D, P=0.003; Fig.4E and P=0.001; Fig.4F). The data demonstrated that, even after normal RT, breast cancer patients with overexpression of IQGAP3 still came to a poor survival.

**Figure 4.**
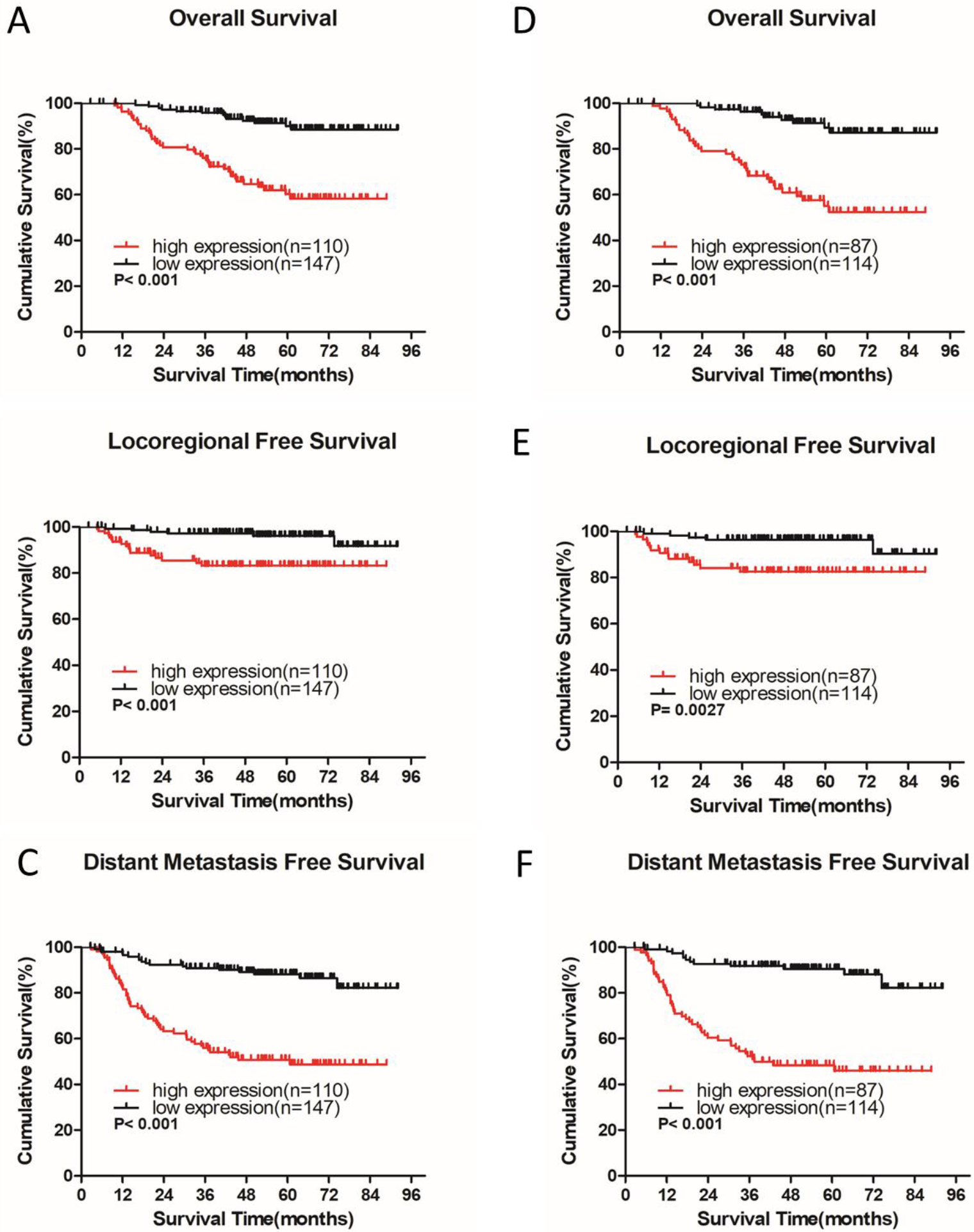
IQGAP3 protein expression is associated with clinical outcomes in the whole cohort of breast cancer cases and in the radiotherapy (RT) subgroup. (A, B, C) Kaplan–Meier overall survival (A), locoregional recurrence-free survival (B) and distant metastasis-free survival (C) curves for all 257 patients with breast cancer stratified by high IQGAP3 expression (n = 110) versus low IQGAP3 expression (n = 147). (D, E, F) Kaplan−Meier overall survival (D),locoregional recurrence-free survival (E) and distant metastasis free survival (F) curves for RT subgroup of 201 patients stratified by high IQGAP3 expression (n = 87) versus low IQGAP3 expression (n = 114). P values were calculated using the log-rank test.

### 2.4. IQGAP3 overexpression act as a bad independent prognostic factor for breast cancer clinical outcome

The clinical variables T category, N category, estrogen receptor (ER) status, progesterone receptor (PR) status, and IQGAP3 expression had an obvious association with survival among the analysis of univariate Cox regression. Multivariate survival analysis also indicated IQGAP3 expression was indeed an independent prognostic factor for OS and progression-free survival (PFS; P = 0.003 and P = 0.001, respectively; Table 2) in the whole cohort breast cancer patients (n=257). Furthermore, we conduct a multivariate survival analysis in the RT subgroup (n=201): IQGAP3 expression remained an independent prognostic factor for OS and PFS (P = 0.002 and P = 0.001, respectively; Table 3) in breast cancer patients who had undergone RT.

Above all, taken these results together: IQGAP3 may be a worthy independent prognostic factor for breast cancer treatment outcome, especially for patients who have undergone RT.

**Table 2.**
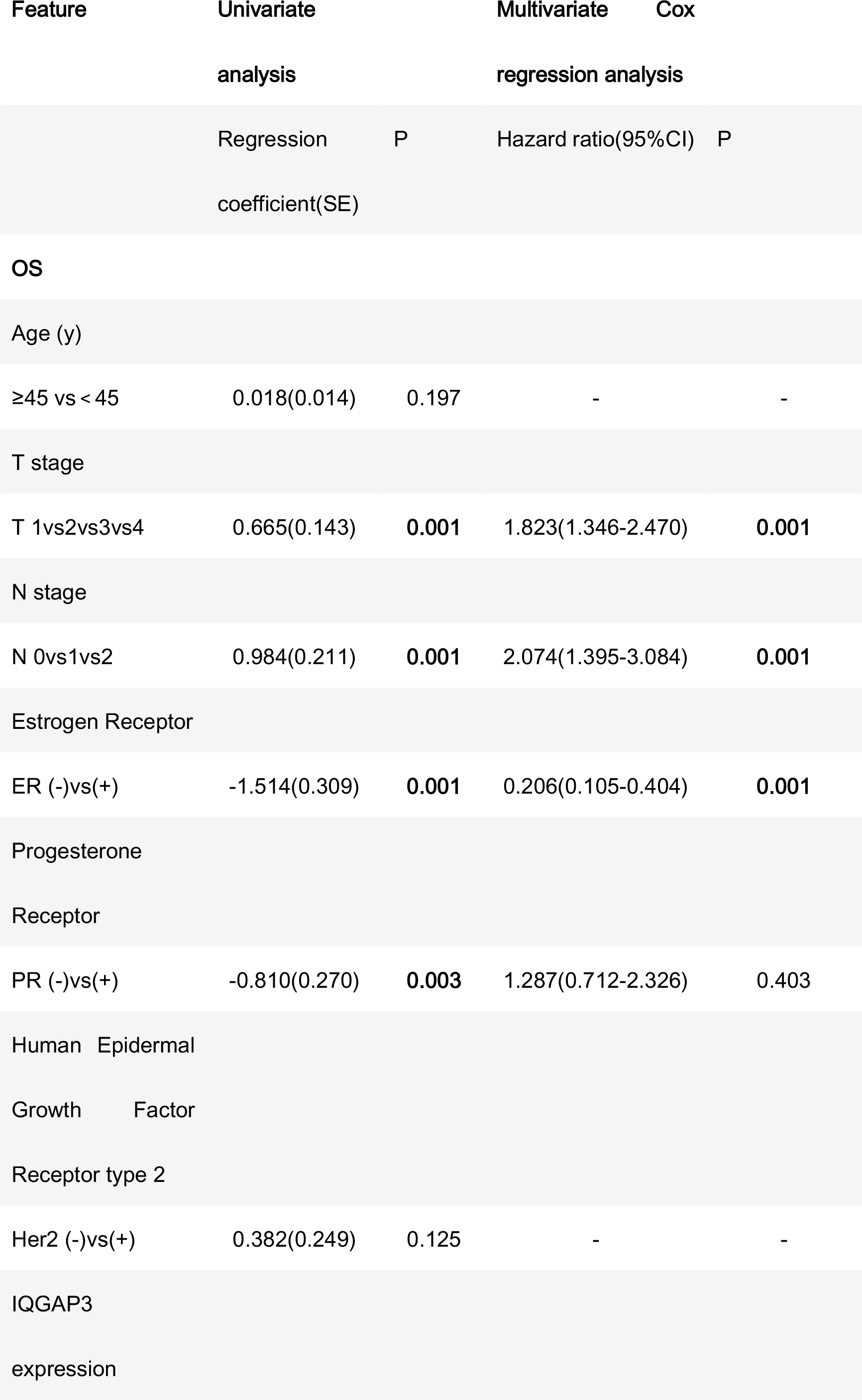
Association of clinicopathological features with overall survival and progression-free survival in breast cancer patients (n=257).

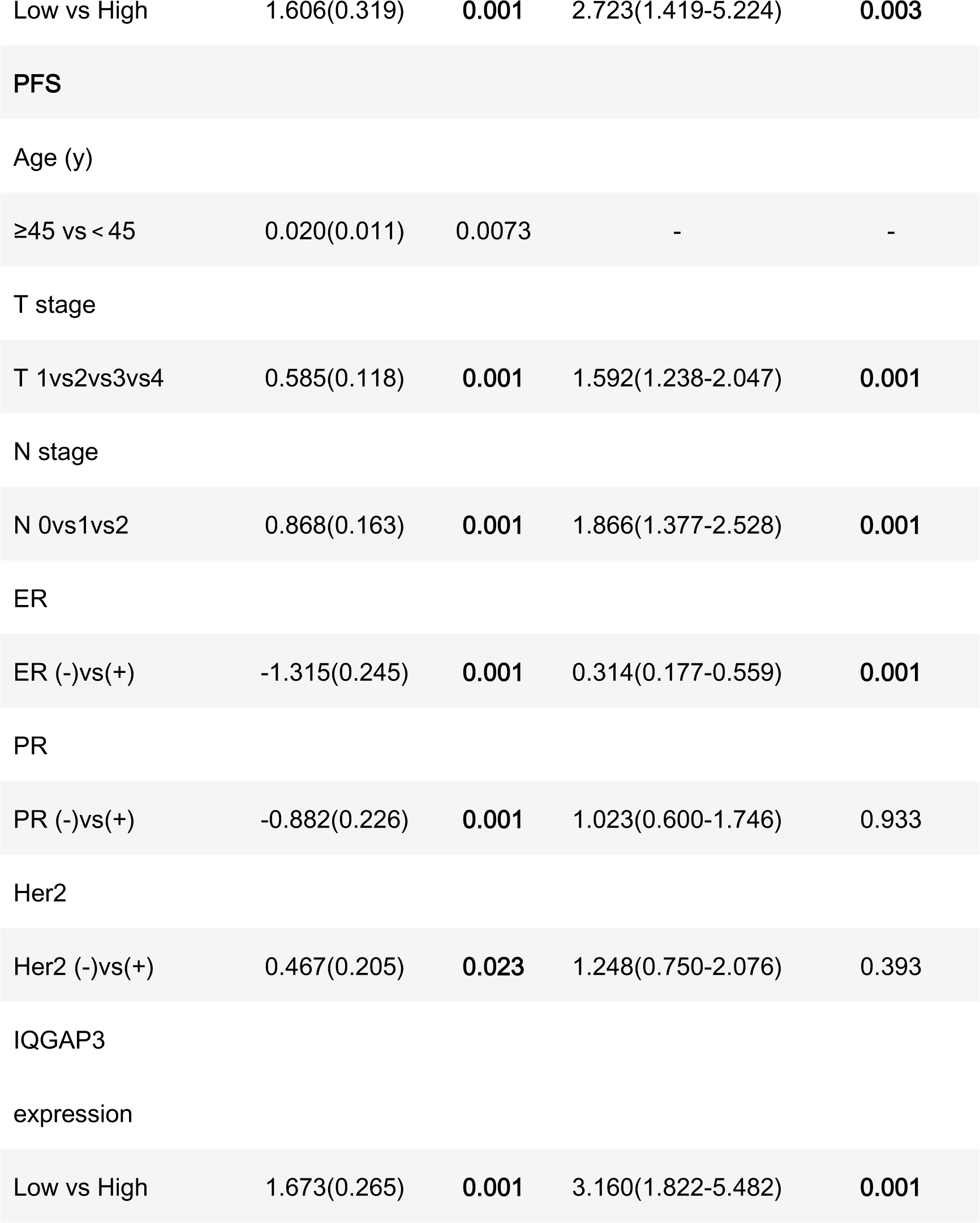

**Table 3.**
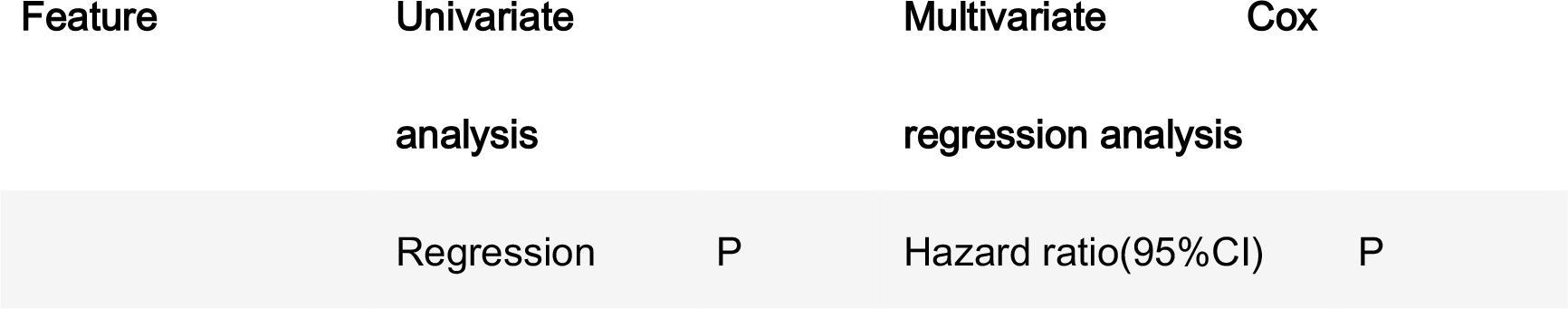
Association of clinicopathological features with overall survival and progression-free survival in breast cancer patients undergoing radiotherapy (n=201).

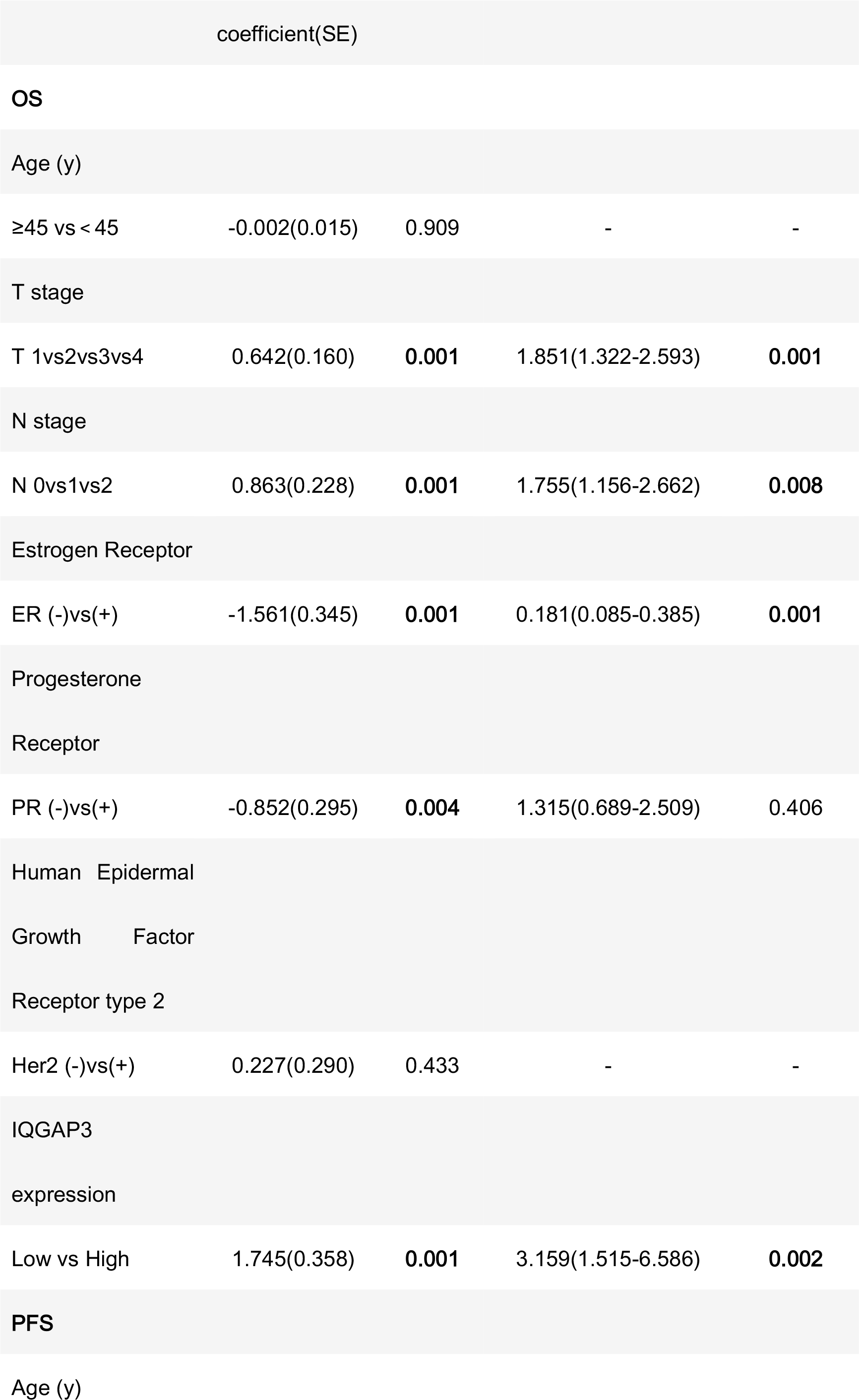

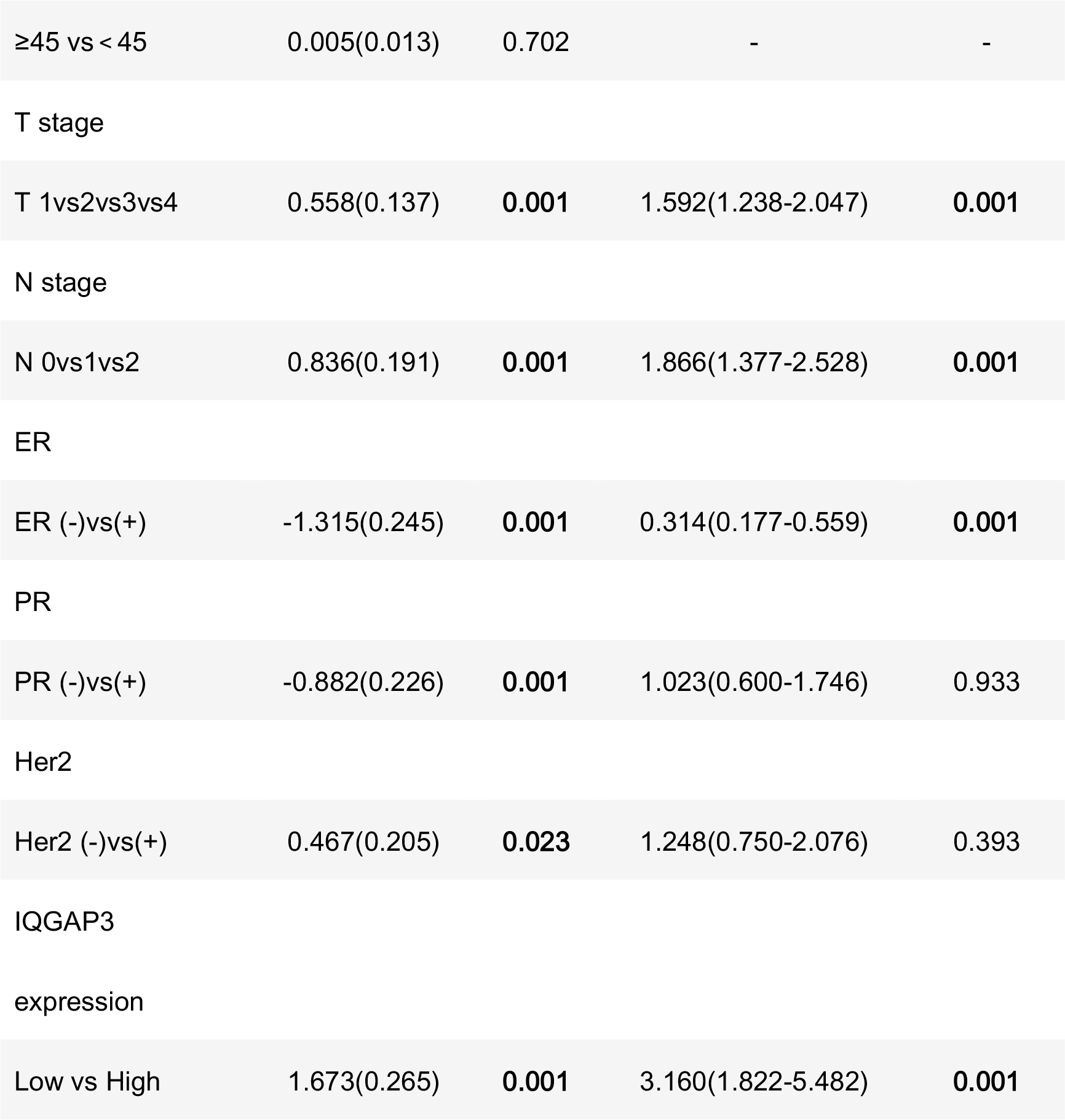

| coefficient(SE) |
| --- |

|  |  |  |  |  |
| --- | --- | --- | --- | --- |
| OS |  |  |  |  |
| Age (y) |  |  |  |  |
| ≥45 vs < 45 | -0.002(0.015) | 0.909 | - | - |
| T stage |  |  |  |  |
| T 1vs2vs3vs4 | 0.642(0.160) | **0.001** | 1.851(1.322-2.593) | **0.001** |
| N stage |  |  |  |  |
| N 0vs1vs2 | 0.863(0.228) | **0.001** | 1.755(1.156-2.662) | **0.008** |
| Estrogen Receptor |  |  |  |  |
| ER (-)vs(+) | -1.561(0.345) | **0.001** | 0.181(0.085-0.385) | **0.001** |
| Progesterone |  |  |  |  |
| Receptor |  |  |  |  |
| PR (-)vs(+) | -0.852(0.295) | **0.004** | 1.315(0.689-2.509) | 0.406 |
| Human Epidermal |  |  |  |  |
| Growth Factor |  |  |  |  |
| Receptor type 2 |  |  |  |  |
| Her2 (-)vs(+) | 0.227(0.290) | 0.433 | - | - |
| IQGAP3 |  |  |  |  |
| expression |  |  |  |  |
| Low vs High | 1.745(0.358) | **0.001** | 3.159(1.515-6.586) | **0.002** |
| PFS |  |  |  |  |
| Age (y) |  |  |  |  |

|  |  |  |  |  |
| --- | --- | --- | --- | --- |
| ≥45 vs < 45 | 0.005(0.013) | 0.702 | - | - |
| T stage |  |  |  |  |
| T 1vs2vs3vs4 | 0.558(0.137) | 0.001 | 1.592(1.238-2.047) | 0.001 |
| N stage |  |  |  |  |
| N 0vs1vs2 | 0.836(0.191) | 0.001 | 1.866(1.377-2.528) | 0.001 |
| ER |  |  |  |  |
| ER (-)vs(+) | -1.315(0.245) | 0.001 | 0.314(0.177-0.559) | 0.001 |
| PR |  |  |  |  |
| PR (-)vs(+) | -0.882(0.226) | 0.001 | 1.023(0.600-1.746) | 0.933 |
| Her2 |  |  |  |  |
| Her2 (-)vs(+) | 0.467(0.205) | 0.023 | 1.248(0.750-2.076) | 0.393 |
| IQGAP3<br>expression |  |  |  |  |
| Low vs High | 1.673(0.265) | 0.001 | 3.160(1.822-5.482) | 0.001 |

### 2.5. IQGAP3 overexpression was significantly correlated with radioresistance in breast cancer

Breast cancer patients who received RT were divided according to their response to treatment into a radioresistant group (who showed disease progression via locoregional recurrence) and a radiosensitive group (without disease progression).

We first analyzed the public microarray TCGA data of breast cancer and found IQGAP3 was overexpressed in radioresistant samples(n=115) compared to radiosensitive samples (n=600; Fig.1B).

To confirm whether IQGAP3 is overexpressed in radioresistant breast cancer patients, we also examined IQGAP3 expression in 159 post-RT patients (radioresistant group n=19; radiosensitive group n=140) using IHC. We demonstrated IQGAP3 was overexpressed in radioresistant breast cancer patients compared to radiosensitive patients (Fig.5).

**Figure 5.**
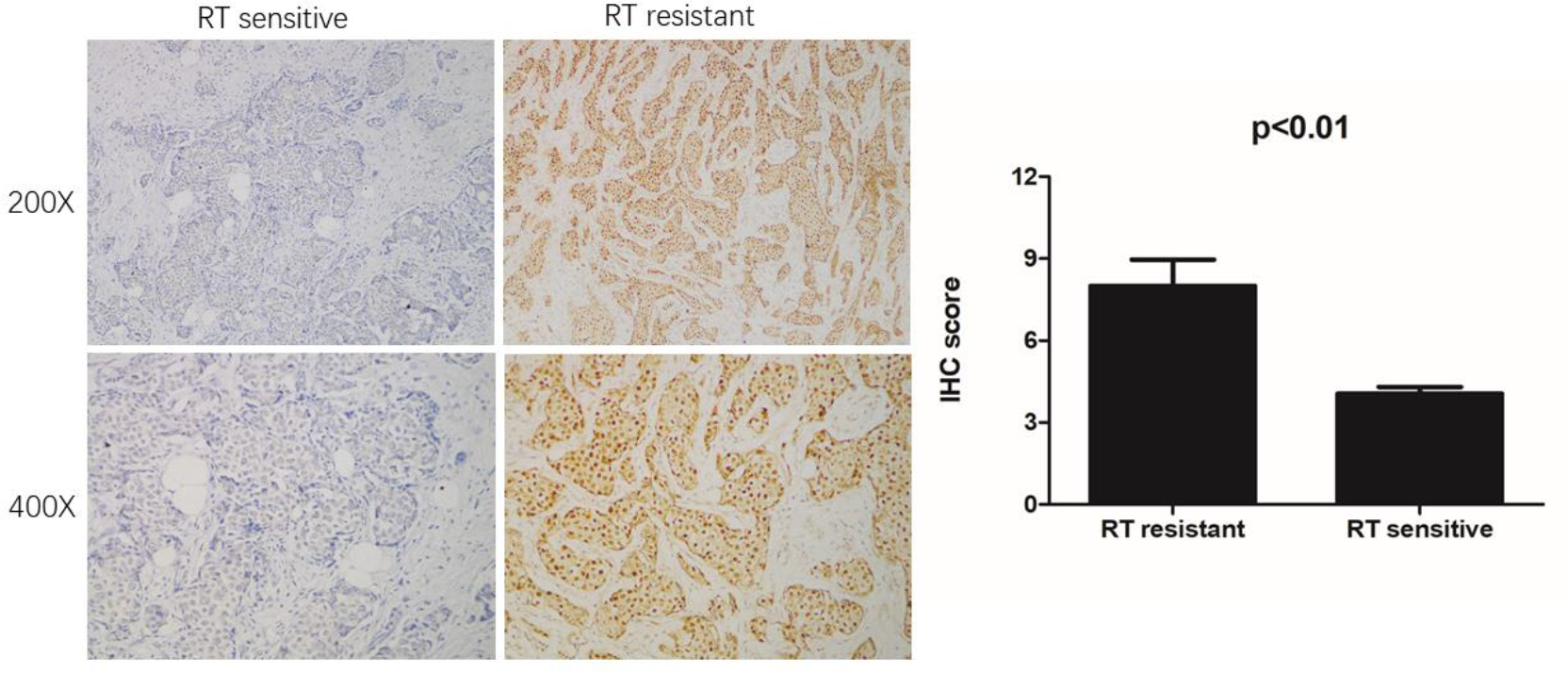
Immunohistochemical (IHC) detection of IQGAP3 expression in paraffin-embedded breast cancer tissues in the radiotherapy (RT) sensitive and RT resistant subgroups. Left panel: Representative images of IHC staining for IQGAP3 in the RT resistant and RT sensitive tissues. Right panel: Average IHC score of IQGAP3 in the RT resistant (n=19) and RT sensitive tissues (n=140).

### 2.6. IQGAP3 overexpression was an independent prognosis factor for radiation therapy outcome in breast cancer

Through further analysis among subgroup of the 159 post-RT cases, we discovered IQGAP3 overexpression was correlated with an obviously shorter radioresistance-free survival (RRFS). Univariate Cox regression analyses showed that IQGAP3 expression, T stage, N stage, ER, and PR were significant prognostic factors for RT outcome (P=0.012, P=0.005, P=0.005, P=0.001 and P=0.001; Table 4). IQGAP3 overexpression remaining an independent prognostic factor for shorter RRFS among multivariate analysis (HR: 3.321; 95% CI: 1.135–9.716; P=0.028).

**Table 4.**
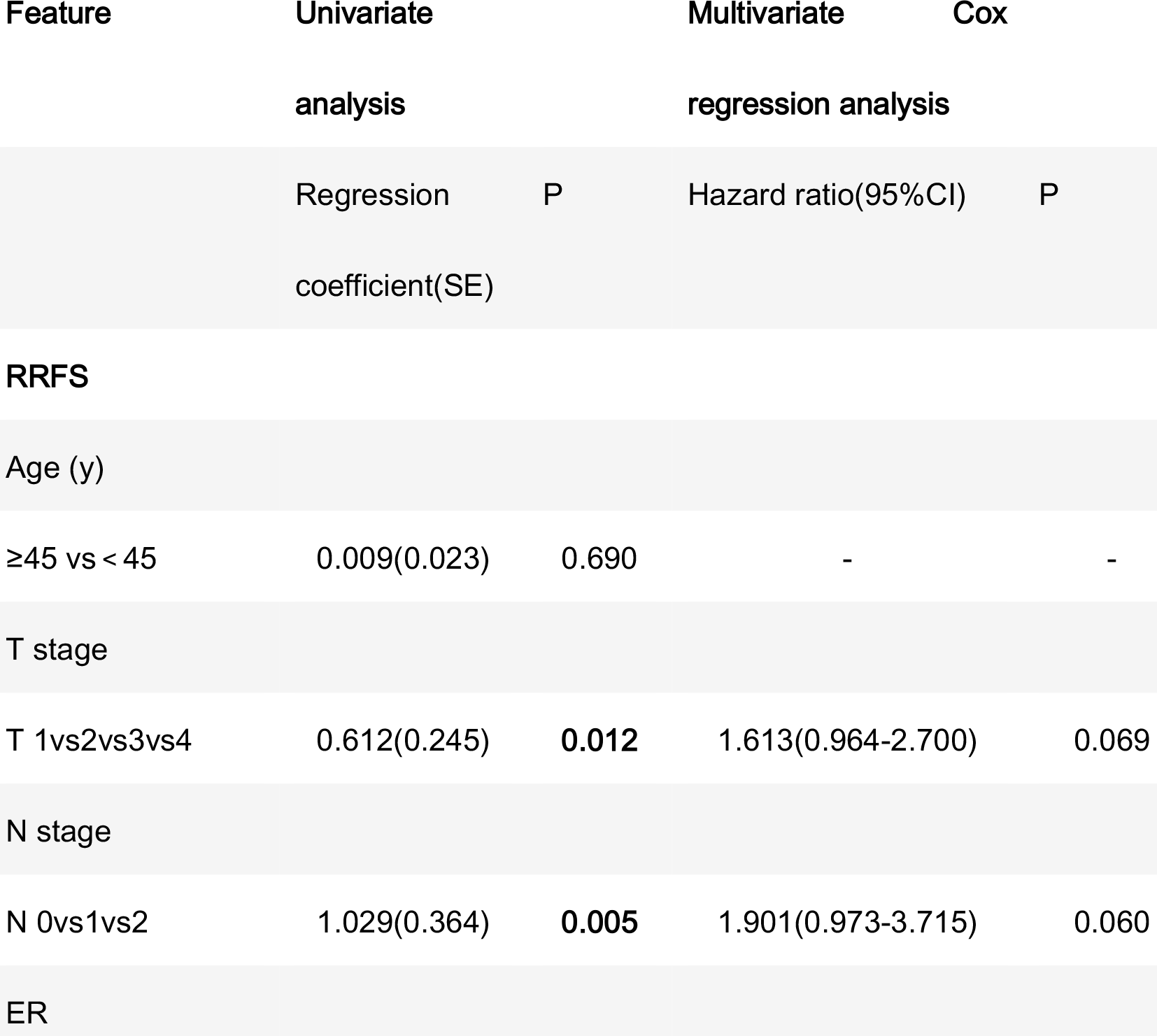
Association of clinicopathological features with radioresistance-free survival in breast cancer cases resistant to radiation therapy (n=159).

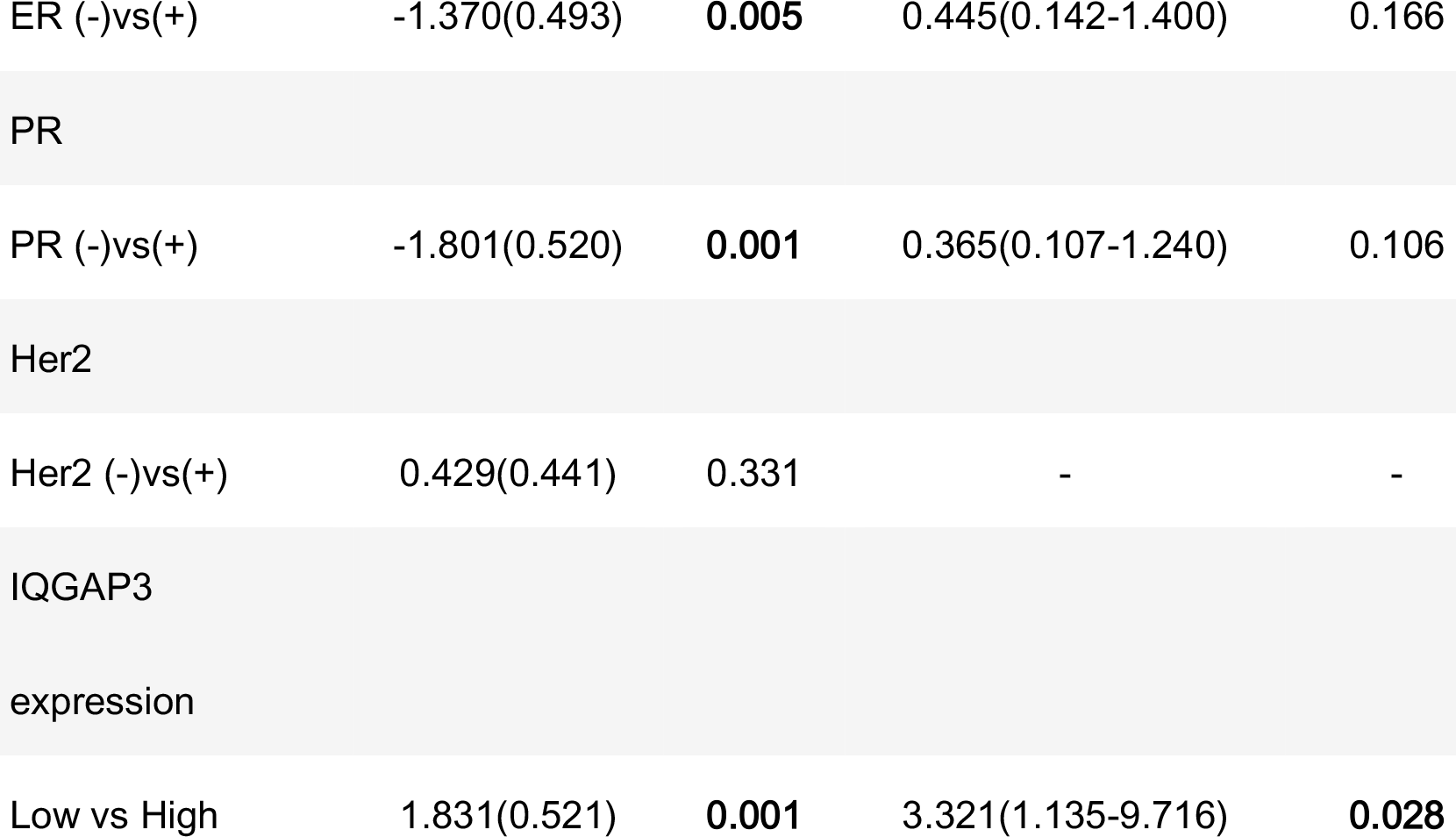

## 3. Discussion

A growing number of evidence reveals that IQGAP3 overexpression in various tumors: for example, melanoma, pancreatic cancer, gastric cancer, bladder cancer, hepatocellular carcinoma, and breast cancer[8-12]. Furthermore, recent study providing significant clue reveals that IQGAP3 overexpression accelerates cell proliferation and invasion in several tumors, showing that IQGAP3 plays a role in the progression of cancer[10,12,13]. However, its specific significance in breast cancer remains vague until now. To our knowledge, we are first to elucidate the tight association between IQGAP3 expression and disease prognosis and RT sensitivity in breast cancer.

IQGAP3 may promote and accelerate cancer development. For example, knocking down of IQGAP3 can inhibit proliferation and ERK activity in cultured epithelial cells[7]; IQGAP3 may help HCC screening and diagnosis by acting as a novel supplementary biomarker [14]; IQGAP3 can active EGFR–ERK signaling and thus promote the metastasis of lung cancer cells[10]. Cell proliferation, migration and invasion, and induced cell apoptosis can be inhibited by knocking down IQGAP3 expression in two pancreatic cancer cell lines BXPC-3 and SW1990[8]. Also, silencing IQGAP3 can inhibit the proliferation, motility, and invasion of breast cancer cell lines[12]. Consistent with this study involving breast cancer cell lines, the present research provides evidence that IQGAP3 expression may have important clinical significance in breast cancer.

We confirmed IQGAP3 was overexpressed both mRNA level (transcriptionally) and protein level (translationally) in breast cancer cell lines and human tumor samples compared to non-cancerous breast epithelial cells and tissues. IQGAP3 overexpression had significant correlation with the following characteristics: gender, clinical stage, T category, N category, vital status, and distant metastasis. Moreover, IQGAP3 overexpression patients were more likely to exhibit locoregional recurrence and distant metastasis, which strongly indicating IQGAP3 protein expression promotes the progression of breast cancer. Also, there is a significant association between high IQGAP3 expression and poorer 5-year OS, LRFS and DMFS in both the entire cohort and the RT-treated subgroup. These above evidence forcefully suggest that IQGAP3 plays a contributive role in the development and progression of breast cancer.

Breast cancer recurrence ranges from 10 to 41%, resting with T status, N status, and tumor grade[15]. Compared to other thoracic tumors, innate or acquired radioresistance leads to locoregional recurrence, and results in treatment failure; therefore, radioresistance represents a formidable clinical problem in the systematic treatment regimen of breast cancer. Although, it is of great importance to identify patients who may resistant to RT before undergoing RT in breast cancer. A specific reliable biomarker could optimize RT has not been established. We found IQGAP3 was strongly positively associated with radioresistance, and high IQGAP3 protein expression had a significant correlation with shorter LRFS and OS even after RT treatment. Therefore, IQGAP3 may be a valuable biomarker to identify specific patients who need more aggressive RT therapeutic regimen (such as a higher dose of radiation) to reduce locoregional recurrence and improve survival. Moreover, by multivariate analysis, we confirmed IQGAP3 was an independent prognostic factor for RRFS in RT sensitive analysis subgroup breast cancer cases. In conclusion, IQGAP3 can act as a reliable novel predictive biomarker of radioresistance and poor survival in breast cancer patients following RT. Therefore, more radical RT can be adopted for patients with high IQGAP3 expression.

Tumor cellular exposure to radiation results in damage to DNA and other cellular structures, which then triggers a complex cascade of downstream response pathways in both the nucleus and cytoplasm, including DNA repair, cell cycle modulation, reactive oxygen species defense, cytokine production, and apoptosis[16]. In certain tumour cell subpopulations, these pathway can be innately biased towards a radioresistant, pro-survival phenotype(i.e., a phenotype with accelerated cell cycle arrest, reduced proliferation, more efficient or prolonged DNA repair, or dampened apoptotic signaling)[17-19]. Indeed, IQGAP3 was found bind to the Ras protein[7], which plays a role in cell cycle arrest, DNA repair, proliferation, and anti-apoptosis in human cancers[20]. Studies have also shown that IQGAP3 can regulate kinds of signaling pathways and cellular functions[21], including mitogen-activated protein kinase (MAPK) signaling, Ca2+/calmodulin signaling, cell–cell adhesion, β-catenin-mediated transcription, and microbial invasion.

To further examine the mechanism of IQGAP3 in the development of breast cancer radioresistance, we performed gene set enrichment analysis (GSEA), using GSEA 2.0.9 (http://software.broadinstitute.org/gsea/index.jsp). We found IQGAP3 expression is positively correlated with DNA repair gene signatures (HALLMARK_DNA_REPAIR) and phosphatidylinositol-4,5-bisphosphate 3-kinase (PI3K) signaling-activated gene signatures (HALLMARK_PI3K_AKT_MTOR_SIGNALING) in TCGA published gene expression profiles (Fig.6). The PI3K signaling pathway, is regulated by Ras, i.e, the direct binding of Ras and the catalytic subunit p110 can directly activate PI3K. This signailing pathway may contribute to the repair, regrowth, redistribution and reoxidation of cells after RT. As evidence, Fan et al. demonstrated that increased expression of PI3K in the breast cancer cell line MDA-DB-453 after RT, not only protects cells from apoptosis, but also significantly enhances their DNA repair ability[22]. In addition, reducing PI3K signaling with a PI3K inhibitor (LY294002) after RT can lead to G2/M cell cycle arrest in a breast cell line MCF-7[23]. On the basis of the above mentioned evidence, we assume that high levels of IQGAP3, combined with Ras, may promote radioresistance in breast cancer by mediating the PI3K signaling pathway.

**Figure 6.**
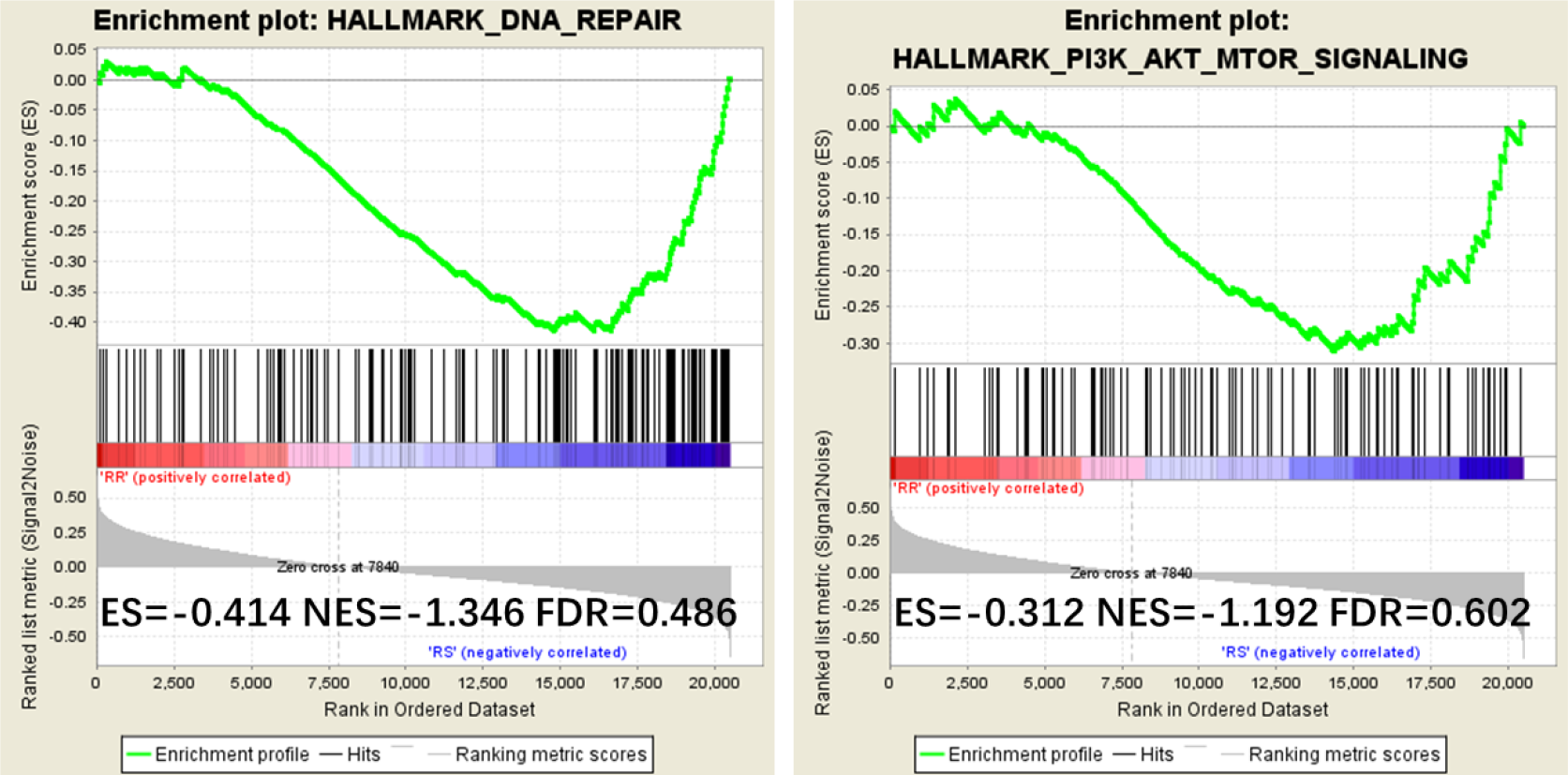
Gene Set Enrichment Analysis (GSEA) plots showing IQGAP3 expression correlates positively with DNA repair gene signatures (HALLMARK_DNA_REPAIR) and PI3K signaling-activated gene signatures (HALLMARK_PI3K_AKT_MTOR_SIGNALING) in published TCGA (the Cancer Genome Atlas) breast invasive carcinoma gene expression profiles.

Although many PI3K inhibitors are currently undergoing investigation in clinical trials, CAL-101 was the first PI3K inhibitor to be approved by the US Food and Drug Administration and the European Medicines Agency for the treatment of different types of leukemia in 2014[24,25]. Therefore, we propose a clinical trial of the use of a PI3K inhibitor as a radiosensitizer should be considered for breast cancer patients with high expression of IQGAP3; however, deeper researches are needed to confirm these hypotheses.

Limitations of our study: first, it was a retrospective study; second, the cohort size was not sufficiently large; and third, some more experiments are need to reveal the precise mechanism of action of IQGAP3 in breast cancer progression and radioresistance.

## 4. Materials and Methods

### 4.1. Microarray data

We performed analyses on the Cancer Genome Atlas (TCGA) data for Breast Invasive Carcinoma-BRCA. Data were acquired from the TCGA data portal https://portal.gdc.cancer.gov/projects/TCGA-BRCA.

### 4.2. Cell lines

Breast cancer cell lines, including MCF-10A, ZR-75-1, SK-BR-3, MDA-MB-468, MDA-MB-453, MCF-7, BT-474, MDA-MB-231,BT-549,HCC1937,SUM159PT,Hs-578T and ZR-75-30 were cultured in DMEM medium (Gibco, Grand Island, NY) supplemented with 10% fetal bovine serum (FBS; HyClone, Logan, UT).

### 4.3. Patients and tissue specimens

In this study, 257 paraffin-embedded breast cancer tissue samples with histopathologically and clinically diagnosed breast cancer were collected from 2006 to 2008 at Sun Yat-sen University Cancer Center (SYSUCC), Guangzhou, China. In addition, six paired breast cancer and adjacent normal tissues were collected from patients who had undergone surgery from 2017 to 2018 at our center. All patients received surgery and 201 (78.2%) received RT after surgery. Clinicopathological classification and staging were determined according to the criteria of the American Joint Committee on Cancer (AJCC). This study was approved by the Clinical Research Ethics Committee of SYSUCC, and written informed consent was obtained from each patient.

### 4.4. PCR

PCR was operated according to the described standard methods [26]. All primers were designed using Primer Express version 2.0 software (Applied Biosystems, Foster City, CA, USA). The primers used were as follows: IQGAP3 forward:5-ATGAGCAGAGGCGGCAGAAT-3, reverse:5-GAACCACGGAGGGTGCAAAA-3; GAPDH forward: 5-GTCTCCTCTGACTTCAACAGCG-3, reverse: 5-ACCACCCTGTTGCTGTAGCCAA-3.

### 4.5. Western blot

Protein Western blotting was carried out according to standard methods as described previously[27], by using anti-IQGAP3 antibody (Abcam, Cambridge, MA). The membranes were stripped and re-probed with an anti-α-tubulin antibody (Sigma, Saint Louis, MI) as a loading control.

### 4.6. Immunohistochemistry

Immunohistochemistry (IHC) and quantification of IQGAP3 expression were performed by two independent pathologists, as previously described[28], by using an anti-IQGAP3 antibody(1:1000; Sigma, Saint Louis, MI). The percentage of cancer cells was scored as 1 (<10%), 2 (10–50%), 3 (50–75%), or 4 (> 75%); and the staining intensity sorted into four grades: 0 (no staining), 1 (weak staining, light yellow), 2(moderate staining, yellow brown), and 3 (strong staining, brown). The scores for the staining intensity and proportion were multiplied. The best cutoff value for IQGAP3 was defined by receiver operating curve(ROC) analysis: a staining score≥6 was classified high expression and score≤4, as low IQGAP3 expression.

### 4.7. Statistical analysis

All statistical analyses were conducted using SPSS (version 20.0; IBM Corporation, Armonk, NY, USA). The survival curves were curved by the GraphPad Prism 6.0 Software. Using the Pearson’s χ^2^ tests or Fisher exact tests to analyze the associations between IQGAP3 expression and clinicopathological features. Using the Kaplan-Meier method and compared via the log-rank test to calculate and curve survival curves. Using Cox’s proportional hazards model to perform multivariate analysis. A two sided P-values < 0.05 were considered statistically significant.

## 5. Conclusions

We are first to report IQGAP3 is overexpressed in breast cancer and its correlation with clinicpathological features. IQGAP3 overexpression was correlated with radioresistance and significantly poorer prognosis. IQGAP3 may be a reliable novel biomarker to provide personalized prognostication and identify patients who may benefit from more aggressive RT treatment for improving the survival of breast cancer patients.

## Authors’ contributions

Conceptualization: LG HXL. Methodology: XH ZQL CL. Software: XH ZQL CL. Validation: ZYH. Formal analysis: XH HXL. Investigation: XQS WW WWZ. Resources: LG HXL WWZ. Data curation: XH HXL ZJL NG. Writing (original draft preparation): XH ZQL. Writing (review and editing): ZYH WWZ HXL. Visualization: XH ZYH ZQL. Supervision: LBS LG HXL. Project administration: LG HXL. Funding acquisition: LG HXL WWZ.

## Ethical approval

All procedures performed in studies involving human participants were in accordance with the ethical standards of the institutional and/or national research committee and with the 1964 Helsinki declaration and its later amendments or comparable ethical standards. This study was approved by the Clinical Research Ethics Committee of SYSUCC.

## Informed consent

Informed consent was obtained from all individual participants included in the study.

